# High prevalence and diversity of nematodes associated with Europe’s most destructive forest pest, revealed by off-target genomic data

**DOI:** 10.64898/2026.11.22.768399

**Authors:** J. Morales-García, K. H. Nowak, P. Zieliński, P. Łukasik, K. Nadachowska-Brzyska

**Affiliations:** Institute of Environmental Sciences, Faculty of Biology, Jagiellonian University, Gronostajowa 7, 30-387, Kraków, Poland; Doctoral School of Exact and Natural Sciences, Jagiellonian University, ul. Łojasiewicza 11, 30-348, Kraków, Poland

**Keywords:** nematodes, spruce bark beetle, off-target reads, whole-genome resequencing

## Abstract

We know very little about the biotic interactions of most species on our planet, including many organisms that play a crucial role in ecosystem networks. One highly cost-effective way of reconstructing these networks is to analyse the rapidly growing collections of high-throughput sequencing datasets. In this study, we examine a large whole-genome resequencing dataset for the spruce bark beetle (*Ips typographus*), one of Europe’s most destructive forest pests, to determine the diversity and distribution of associated organisms that such datasets can reveal. We found that from among metagenomic datasets for 312 adult beetles from across the species’ European range, 69% contained small subunit ribosomal RNA (SSU rRNA) reads from other eukaryotic organisms, most often nematodes. Among 13 rRNA-reconstructed nematode species, *Contortylenchus* sp. and *Micoletzkya* sp. were the most prevalent taxa across the European populations. Frequently, nematode DNA comprised a large proportion of the sequencing libraries, suggesting a significant impact on the host. Mitochondrial genes reconstructed for many of these beetles provided additional phylogenetic resolution, although the reconstruction success varied. Overall, our results demonstrate potential for reconstructing the key biological associations using sequencing data originally generated for a different purpose, highlighting a path towards comprehensive survey of interactions likely to substantially influence the population dynamics of ecologically significant species.

## Introduction

Biodiversity plays a key role in ecosystem functioning, but in many systems, it remains incompletely described. We often lack basic knowledge on species presence in ecosystems and their ecological roles. Further, even for species that have been the focus of extensive research, the diversity of interactions with other organisms is rarely fully characterized and understood (Ezenwa et al., 2016; Holt et al., 2024). These hidden relationships with parasites, commensals, and mutualists representing bacteria, protists, fungi, small invertebrates, and other taxonomic groups have the potential to influence various aspects of host biology, including survival, reproduction, dispersal, and behaviour (Rea & Irwin, 1994; Susoy & Herrmann, 2014; Ezenwa et al., 2016; Stephenson & Adelman, 2022; Campusano et al., 2026). Consequently, understanding these interactions can help explain how associated taxa influence host population dynamics.

The rapid development of high-throughput sequencing technologies has facilitated the generation of immense volume of genomic data for wild organisms, enabling researchers to address a wide range of ecological and evolutionary hypotheses (Satam et al., 2023; Theissinger et al., 2023; Combrink et al., 2025). Yet such genomic datasets, typically generated with a specific research question in mind, may contain a wealth of additional information beyond the original scope of the study (Campusano et al., 2026). For example, in addition to the targeted organism, they may contain the DNA from a wider array of physically associated taxa: microbial symbionts, parasites, commensals and pathogens (Ghanavi et al., 2021; Campusano et al., 2026). Therefore, genomic datasets prepared from whole wild-caught individuals (e.g. whole insects) should be considered as metagenomes, with a strong potential to inform us of the biological interactions of focal taxa. Reads corresponding to these non-target organisms, often referred to as “off-target”, are typically filtered and discarded during analyses focused on the host genome.

However, researchers increasingly realize the potential of such off-target data. For example, metagenomic datasets for diverse British insects have yielded a large collection of complete genomes of their symbiont *Wolbachia* (Vancaester & Blaxter, 2023) and microsporidial parasites (Khalaf et al., 2025), while bycatch reads from genomic and target enrichment dataset–from *Dendroctonus* beetles have provided insight into genome and population structure of their associated *Contortylenchus* nematodes (Campbell et al., 2026; Campusano et al., 2026). It is clear that as whole-genome sequencing and resequencing datasets continue to accumulate, they could offer an invaluable window into biotic interactions of target taxa without additional sampling or sequencing efforts.

The spruce bark beetle, *Ips typographus*, is one of the most destructive forest pests in Europe (Christiansen & Bakke, 1988; Wermelinger, 2004a). This bark-boring insect, widely distributed across Eurasia, primarily colonizes Norway spruce (*Picea abies*). The beetles excavate galleries in the phloem and inoculate ophiostomatoid fungi into the sapwood, and together these processes disrupt nutrient and water transport (Wermelinger, 2004b; Biedermann et al., 2019). Mass beetle attacks can overwhelm host defences and kill even vigorous trees, and their large-scale outbreaks have caused substantial economic losses and severe environmental impacts in Norway spruce forests across Europe (Seidl et al., 2016; Hlásny et al., 2021). Although the effects of abiotic factors on bark beetle outbreaks – including drivers such as temperature, drought, or large-scale windthrow events – have been extensively studied (e.g., Wermelinger, 2004b; Seidl et al., 2016), the role of biotic interactions remains comparatively poorly explored despite their strong potential to influence outbreak dynamics (Biedermann et al., 2019). For example, the extent to which natural enemies, including predators, parasites or parasitoids, pathogens (e.g. entomopathogenic fungi or viruses), and phoretic organisms (e.g. mites) affect spruce bark beetle populations is still largely unknown (Vega & Hofstetter, 2015; Wegensteiner et al., 2015; Biedermann et al., 2019). The availability of whole-genome resequencing data from hundreds of *I. typographus* individuals sampled across Europe (Mykhailenko et al., 2024; Zieliński et al., 2026) provided an opportunity to investigate such host-associated organisms across a large portion of this species’ geographic range.

Here, we asked whether an extensive *I. typographus* genomic resequencing dataset, spanning 312 individuals from 23 populations, can be used to characterise the diversity, prevalence, and geographic structure of organisms associated with this important forest pest. First, we tested whether reads unmapped to the bark beetle genome (off-target reads) contain detectable signatures of host-associated taxa, and whether their abundance is linked to reduced mapping rates to the host genome. Second, we assessed the taxonomic diversity of organisms detected in beetle genomic datasets and, for the most abundant group – nematodes – characterized their community composition. Third, we evaluated the geographic patterns in nematode distribution and prevalence. Finally, we assessed whether genomic regions offering and greater phylogenetic resolution than rRNA markers could be reliably reconstructed for the predominant nematode lineages associated with *I. typographus*.

## Materials and Methods

### Study organisms

The study encompassed 312 adult individuals of spruce bark beetle (*I. typographus*), representing 23 different populations distributed across European species range (11-15 per population; Table S1, Figure S1). These populations were divided into three regions based on the species’ genetic structure (northern, southern and central region; Table S1, Mykhailenko et al., 2024; and Zieliński et al., 2026). Individuals trapped using pheromone traps were either preserved in ethanol or processed alive shortly after collection, depending on the site and logistic constraints. Whole-genome data were generated for the population genomic analyses of *I. typographus*, as described in Mykhailenko et al., 2024; and Zieliński et al., 2026. Briefly, whole-body DNA was extracted from individual beetles using the Wizard Genomic DNA Purification Kit (Promega). Sequencing libraries were prepared using the NEBNext Ultra II FS DNA Library Prep Kit (New England Biolabs) and sequenced using Illumina NovaSeq 6000 platform (2 × 150 bp mode; Swedish National Genomics Infrastructure, SNP&SEQ Technology Platform and Novogene Company) to an average coverage of 8.3 Gb (SD = 2.4 Gb) of data per individual.

### Genomic data processing

For the metagenomic analysis, we used quality-trimmed sequencing reads (FASTQ files; Andrews, 2010) obtained from each individual. Taxonomic profiling was performed using phyloFlash v3.4 (Gruber-Vodicka et al., 2020), which reconstructs and classifies small-subunit (SSU; eukaryotic 18S and bacterial 16S) ribosomal RNA (rRNA) gene sequences directly from metagenomic data. The -almosteverything flag was enabled for each individual sample to include SSU rRNA read extraction, SSU rRNA sequence reconstruction, and taxonomic classification, with resolution set at the species level (-taxlevel 8). To assess the reliability of phyloFlash taxonomic assignments, we compared the corresponding reference sequences against the NCBI nucleotide database using BLASTn (Altschul et al., 1990). In two cases, species-level assignments were inconsistent with BLASTn results and were replaced with genus-level assignments supported by these results (≥97% sequence identity).

For downstream analysis, we considered only taxa for which SSU rRNA sequences could be successfully assembled with phyloFlash, excluding detections based only on unassembled reads to reduce the risk of spurious assignments from short or low-quality fragments or potential low-level cross-contamination through index hopping. To account for differences in sequencing depth and library composition among samples, relative abundance was estimated as the proportion of SSU rRNA reads assigned to a given taxon relative to the total SSU rRNA reads recovered for that individual. This relative metric was used because the variation in overall coverage of the sequencing libraries differed across samples, making direct comparisons of absolute read counts unreliable.

In addition, to assess the prevalence of the most common organisms associated with the spruce bark beetle across its populations, we quantified species occurrence using a presence/absence approach. For each population, we counted the number of beetles in which at least one associate SSU rRNA sequence was successfully assembled, irrespective of the total number of reads. Finally, to assess the robustness of taxonomic assignments, we re-ran phyloFlash on a set of 9 individuals for which duplicate libraries were available (independent libraries prepared and sequenced from the same DNA extraction of an individual; Mykhailenko et al., 2024).

### Prevalence and diversity of associated taxa

The proportion of beetle DNA within individual metagenomes (mapping rate) was assessed using quality trimmed paired-end reads and the *I. typographus* reference genome with Bowtie2 (Langmead & Salzberg, 2012). To test whether lower mapping rates were associated with a higher proportion of SSU rRNA reads assigned to other associated organisms, we performed two complementary analyses. First, to assess whether variation in mapping rates among libraries was associated with differences in the proportion of SSU rRNA reads assigned to associated organisms, we tested for a correlation between mapping rate to the spruce bark beetle genome and the proportion of SSU rRNA reads not assigned to *I. typographus* (non-*Ips* SSU rRNA reads / total SSU rRNA reads). Second, to assess if only the most abundant associate group may explain the observed differences in mapping rates, we tested for a correlation between mapping rate and the within-individual ratio of SSU rRNA reads assigned to this dominant associate group relative to *I. typographus* SSU rRNA reads (dominant associate SSU / *Ips* SSU). Correlations were assessed using Spearman’s rank correlation (ρ) and visualised on a logarithmic scale to highlight substantial variation in the representation of associated taxa among individuals.

To characterise the diversity of organisms associated with *I. typographus* across its geographic range, phyloFlash-assembled SSU rRNA sequences were first taxonomically classified per individual spruce bark beetle library. Species-level assignments were retained only for taxa showing ≥99% SSU rRNA sequence identity in phyloFlash and for which species identification was supported by previous literature on *Ips typographus* associated microorganisms (Muratoğlu et al., 2011; Grucmanová & Holuša, 2013; Cheng et al., 2023). Taxa not meeting these criteria were conservatively reported at the genus level.

The diversity of nematode taxa (the most predominant associate group) was assessed across populations by compiling the list of nematode taxa detected by phyloFlash in each geographic group. To test whether the occurrence of specific nematode taxa differed among geographic groups, we focused on the four most frequently detected species (*Contortylenchus* sp., *Micoletzkya* sp., *Bursaphelenchus* sp. and *Parasitorhabditis obtusa*) and tested for an association between geographic group and taxon presence/absence using contingency table analyses. Pearson’s Chi-squared tests were used when expected cell counts were sufficient (≥5); otherwise, Fisher’s exact tests were used. Additionally, to assess whether the observed geographically structured distribution of nematodes taxa could be explained by chance, we performed permutation tests of nematode occurrence across geographic groups. For each nematode taxon, we randomly shuffled detection and non-detection labels among individual beetles 1000 times, keeping each beetle’s geographic group (N, C, or S) fixed. This preserved the number of beetles sampled in each group and the total number of detections for that taxon, while allowing the number of detections in each group to vary across permutations. P-values were adjusted for multiple testing using false discovery rate (FDR) correction. All statistical analyses were conducted in R (R Core Team 2024).

### Phylogenetic analysis and mitogenomes reconstruction

First, we reconstructed the evolutionary relationships among species based on full-length SSU rRNA in MEGA11 (Tamura et al., 2021). The unique sequences (one per haplotype) were aligned using Muscle implemented in MEGA11 with default parameter settings. The phylogenetic tree was reconstructed using the Maximum Likelihood method assuming the Tamura-Nei model of sequence evolution (Tamura & Nei 1993).

Second, mitochondrial sequences from 312 samples were assembled from paired-end Illumina reads using GetOrganelle v1.7.7.0 (Jin et al., 2020). Each sample was processed independently with get_organelle_from_reads.py using the animal_mt assembly mode, 25 extension rounds (-R 25), and multi-k-mer strategy (21,45,65,85,105). To recruit divergent mitochondrial reads, a custom reference file containing six representative nematode mitogenomes was used via the --genes option (Table S2).

GetOrganelle generated assemblies were classified using BLASTN (blastn: 2.17.0+ Package: blast 2.17.0) against a custom local database consisting of the abovementioned nematode mitogenomes and the spruce bark beetle mitogenome (https://www.ncbi.nlm.nih.gov/nuccore/OK110249.1). The scaffolds that did not produce significant matches against the local reference database were searched against the full NCBI nucleotide database. The classified scaffold sets were annotated independently using MITOS2 v2.1.10 (Bernt et al., 2013) using the metazoan reference dataset and the --best option to retain the highest-confidence annotation for each feature. Finally, protein-coding genes annotated by MITOS2 were extracted and independently aligned. From genes recovered by MITOS2, COX1 and COX2 were used for further phylogenetic comparison.

Phylogenetic relationships among nematodes were assessed separately using SSU rRNA, COX1, and COX2 sequences. For each marker, two datasets were prepared: sequences clustered at 99% nucleotide identity and sequences without clustering, yielding six phylogenetic analyses.

Clustering was performed with a wrapper script running VSEARCH (Rognes et al., 2016) using --cluster_smallmem --usersort --id 0.99 --iddef 2 --strand plus --qmask none. Sequences were ordered by decreasing frequency of identical sequences in the input, with sequence length used to resolve ties. Each cluster was represented by an original seed sequence rather than a consensus.

SSU and COX1 reference sequences were retrieved from GenBank and ENA, selecting up to two representatives per target genus according to taxonomic identification, sequence coverage, and documented insect associations where available. The SSU panel included 15 ingroup references covering ten genera and *Plectus aquatilis* (AY284700.1) as an outgroup. The COX1 panel contained 14 references covering eight genera, obtained from annotated gene records or extracted from mitochondrial genomes. Accession numbers and source information are provided in the Table S3. References were added to the clustered and dereplicated SSU rRNA sequences as well as to the unique COX1 sequences; none were added for COX2.

Each dataset was aligned independently with MUSCLE v3.8.425 (Edgar, 2004) in AliView (Larsson, 2014). Within the alignments, columns containing a nucleotide in only one sequence were removed using a custom script. Trimming of COX1 and COX2 alignments was codon-aware to preserve the reading frame.

Maximum-likelihood trees were inferred using IQ-TREE v3.0.1 (Wong et al., 2026). SSU was analysed without partitioning, with substitution-model selection using ModelFinder (-m MFP; Kalyaanamoorthy et al., 2017). COX1 and COX2 were initially partitioned by codon position, with model selection and partition merging performed using -p codon_partition.txt -m MFP+MERGE. Branch support was assessed with 1000 ultrafast bootstrap replicates and 1000 SH-aLRT replicates (-B 1000 --alrt 1000), enabling additional bootstrap-tree optimization (--bnni). Trees were visualized in iTOL (Letunic & Bork, 2007). SSU trees were rooted using *Plectus aquatilis* (AY284700.1), whereas COX1 and COX2 trees were midpoint-rooted without an outgroup. Trees were exported, displaying UFBoot values below 100% and omitting labels for branches with 100% support.

## Results

### Off-target SSU rRNA profiling reveals abundant biological associates in spruce bark beetle genome resequencing datasets

We observed that mapping rates of *I. typographus* resequencing datasets to the reference genome of the species varied considerably among individuals, with a mean unmapped fraction of 16% (7% - 61%, SD=10%) (Figure 1, top panel). We then used these datasets for SSU rRNA sequence reconstruction to explore whether abundant DNA from other organisms was present, potentially explaining the variation in host mapping rates. SSU rRNA sequences assigned to *I. typographus* were successfully recovered from 95.2% of metagenomic datasets for individual beetles, but at the same time, from 79.2% datasets, we also reconstructed the sequences of diverse other organisms. Based on phyloFlash-reconstructed coverage, the relative abundance of host SSU rRNA varied markedly among individuals, ranging from 0 to 100% (mean: 76.8%) (Figure 1, middle panel).

**Figure 1.**
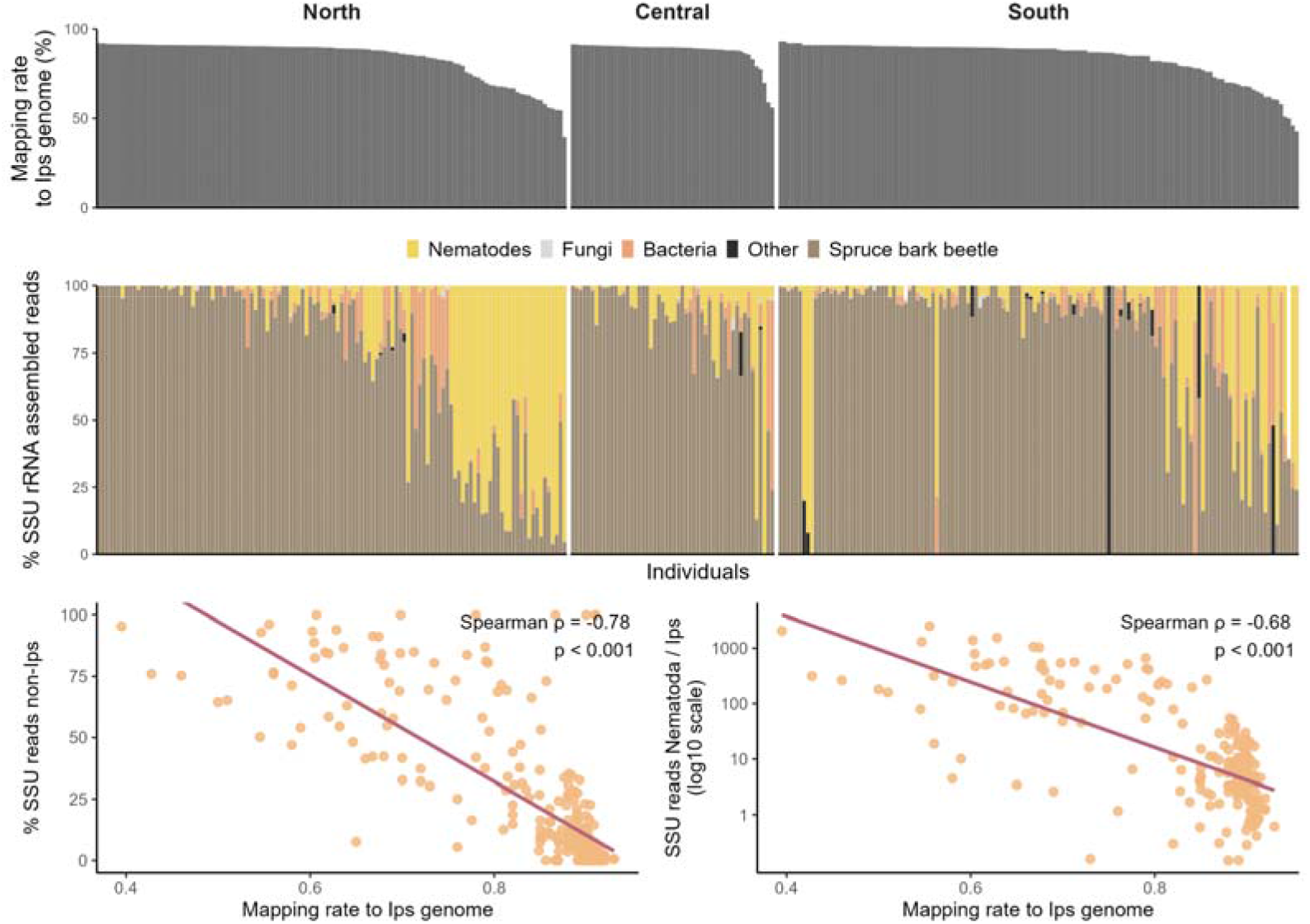
The taxonomic composition of whole-beetle genomic libraries for 312 *Ips typographus* individuals. Top panel: the proportion of metagenomic reads from individual libraries that mapped to the *I. typographus* reference genome. Individuals, represented by distinct bars, are sorted by the collection regions and, within each region, by descending mapping rate to the *I. typographus* genome. Middle panel: the relative abundance of SSU rRNA copies of different organisms across individual metagenomic datasets, based on phyloFlash-estimated coverage of alternative SSU variants. Individuals, represented by columns, are listed in the same order as in the top panel. Bottom panel: correlations between mapping rate to the *I. typographus* genome and the relative abundance of non-host (left plot) and nematode (right plot) SSU rRNA reads. Each point represents an individual sample, and lines indicate linear correlation fits. Spearman’s ρ and p-values are indicated in each panel.

Metagenomic read mapping rates to the *I. typographus* reference genome were negatively correlated with the proportion of non-host SSU rRNA reads (Spearman’s ρ = −0.78, p < 0.001; Figure 1, left bottom panel). Nematodes were the most frequently detected group of associated organisms, and the relationship followed the same pattern when considering nematode SSU rRNA specifically (Spearman’s ρ = −0.68, p < 0.001; Figure 1, right bottom panel). Nematode detection showed 83% concordance between duplicate sequencing libraries (Table S4).

### The taxonomic diversity of beetle-associated organisms

SSU rRNA profiling detected nematodes, bacteria, fungi, mites and apicomplexans in spruce bark beetle datasets (Table 1). Nematodes were the most frequently detected group, occurring in 216 of 312 beetles (69.2%); their diversity and distribution is discussed in detail below. Bacterial rRNA sequences were reconstructed from 146 of 312 beetles (46.8% samples), with *Acinetobacter* sp. being the most prevalent taxon (18% of datasets with SSU detections), followed by *Pseudomonas* spp. (10.3%), *Rahnella* sp. (9.2%), *Erwinia* spp. (7.4%) and *Serratia* spp. (6.7%). Some showed marked geographic differences: *Acinetobacter* sp. occurred in all three groups but was most frequent in central populations (25%), whereas *Pseudomonas* spp. were detected in northern and southern populations but not in the central group. Fungi, mites and apicomplexans were seen less frequently. The prevalence of individual taxa varied markedly among geographic groups, with the full breakdown shown in Supplementary Information - Other Materials.

**Table 1.** Taxonomic groups and taxa detected from off-target SSU rRNA sequences recovered from *Ips typographus* whole-genome resequencing libraries. Prevalence in North, Central, and South regions was calculated as the proportion of individuals in each geographic region in which a given taxon was detected, expressed as a percentage of beetle individuals in which the taxon was detected. Overall prevalence was calculated analogously across all 312 individuals included in the study. Rows are ordered by decreasing overall prevalence. The functional categories are assigned to broad taxonomic clades that are likely associated with beetles in similar manner. Only taxa with an overall prevalence ≥ 1% are shown; the complete list of detected taxa is provided in Supplementary Information – Other Materials.

| Category | Order | Taxon | North (%) | Central (%) | South (%) | Overall (%) |
|---|---|---|---|---|---|---|
| **Nematodes** | Diplogasterida | *Micoletzkya* sp. | 43.3 | 73.1 | 69.9 | 60 |
|  |  | *Diplogasteroides halleri* | 1.7 | 0 | 9 | 4.6 |
|  | Rhabditida | *Parasitorhabditis obtusa* | 4.2 | 5.8 | 7.5 | 5.9 |
|  |  | *Odontopharynx* sp. | 0 | 0 | 6 | 2.6 |
|  | Tylenchida | *Contortylenchus* sp. | 35 | 7.7 | 12 | 20.3 |
|  |  | *Bursaphelenchus* sp. | 3.3 | 11.5 | 3.8 | 4.9 |
|  |  | *Cryptaphelenchus* sp. | 4.2 | 5.8 | 0.8 | 3 |
|  |  | *Parasitylenchus* sp. | 0.8 | 0 | 4.5 | 2.3 |
|  |  | *Ektaphelenchus* sp. | 0 | 0 | 3.8 | 1.6 |
|  |  | *Deladenus* sp. | 2.5 | 0 | 1.5 | 1.6 |
| **Bacteria** | Burkholderiales | *Janthinobacterium* sp. | 0 | 0 | 6.8 | 3 |
|  | Enterobacterales | *Rahnella* sp. | 14.2 | 0 | 8.3 | 9.2 |
|  |  | *Serratia* spp. | 7.5 | 11.5 | 4.5 | 6.7 |
|  |  | *Lelliottia* sp. | 0 | 7.7 | 1.5 | 2 |
|  |  | *Serratia liquefaciens* | 2.5 | 0 | 2.3 | 2 |
|  |  | *Rouxiella* sp. | 1.7 | 0 | 1.5 | 1.3 |
|  |  | *Erwinia* spp. | 4.2 | 1.9 | 12.8 | 7.4 |
|  |  | *Hafnia* spp. | 0 | 7.7 | 1.5 | 1.9 |
|  | Entomoplasmatales | *Spiroplasma* sp. | 1.7 | 0 | 12 | 5.9 |
|  | Flavobacteriales | *Wautersiella* sp. | 0 | 7.7 | 0 | 1.3 |
|  | Lactobacillales | *Lactococcus* sp. | 0 | 15.4 | 2.3 | 3.6 |
|  |  | *Enterococcus* sp. | 0 | 0 | 2.3 | 1 |
|  | Pseudomonadales | *Acinetobacter* sp. | 12.5 | 25 | 20.3 | 18 |
|  |  | *Pseudomonas* spp. | 9.2 | 0 | 15.8 | 10.3 |
|  | Rickettsiales | *Wolbachia* | 5 | 0 | 2.3 | 3 |
|  | Xanthomonadales | *Stenotrophomonas* sp. | 0 | 5.8 | 0 | 1 |
| **Fungi** | Mortierellales | *Mortierella* sp. | 0 | 0 | 5.3 | 2.3 |
|  | Saccharomycetales | *Yarrowia* sp. | 0 | 11.5 | 0 | 2 |
|  |  | *Wickerhamomyces bisporus* | 0 | 0 | 2.3 | 1 |
| **Acari** | Mesostigmata | *Trichouropoda* sp. | 0.8 | 0 | 3 | 1.6 |
|  | Unresolved | Unresolved sp. | 2.5 | 5.8 | 10.5 | 6.6 |
| **Apicomplexa** | Neogregarinorida | *Mattesia* sp. | 0 | 0 | 2.3 | 1 |

### Nematode distribution across geographic groups

Thirteen nematode taxa, differing substantially in prevalence and distribution, were detected across the complete SSU dataset (Table 1). *Micoletzkya* sp. was the most prevalent, occurring in 60% of datasets with SSU detections, more often in central (73.1%) and southern populations (69.9%) than in the North (43.3%). *Contortylenchus* sp. was the second most prevalent taxon (20.3% overall) but showed the opposite geographic pattern: it occurred in 35% of northern datasets, compared to 7.7% of central and 12% of southern datasets (Table 1, Figure 2, Table S5). The prevalence of both nematode species varied significantly among geographic regions (*Micoletzkya* sp.: χ² = 18.27; *Contortylenchus* sp.: χ² = 28.45; FDR-corrected p < 0.001).

**Figure 2.**
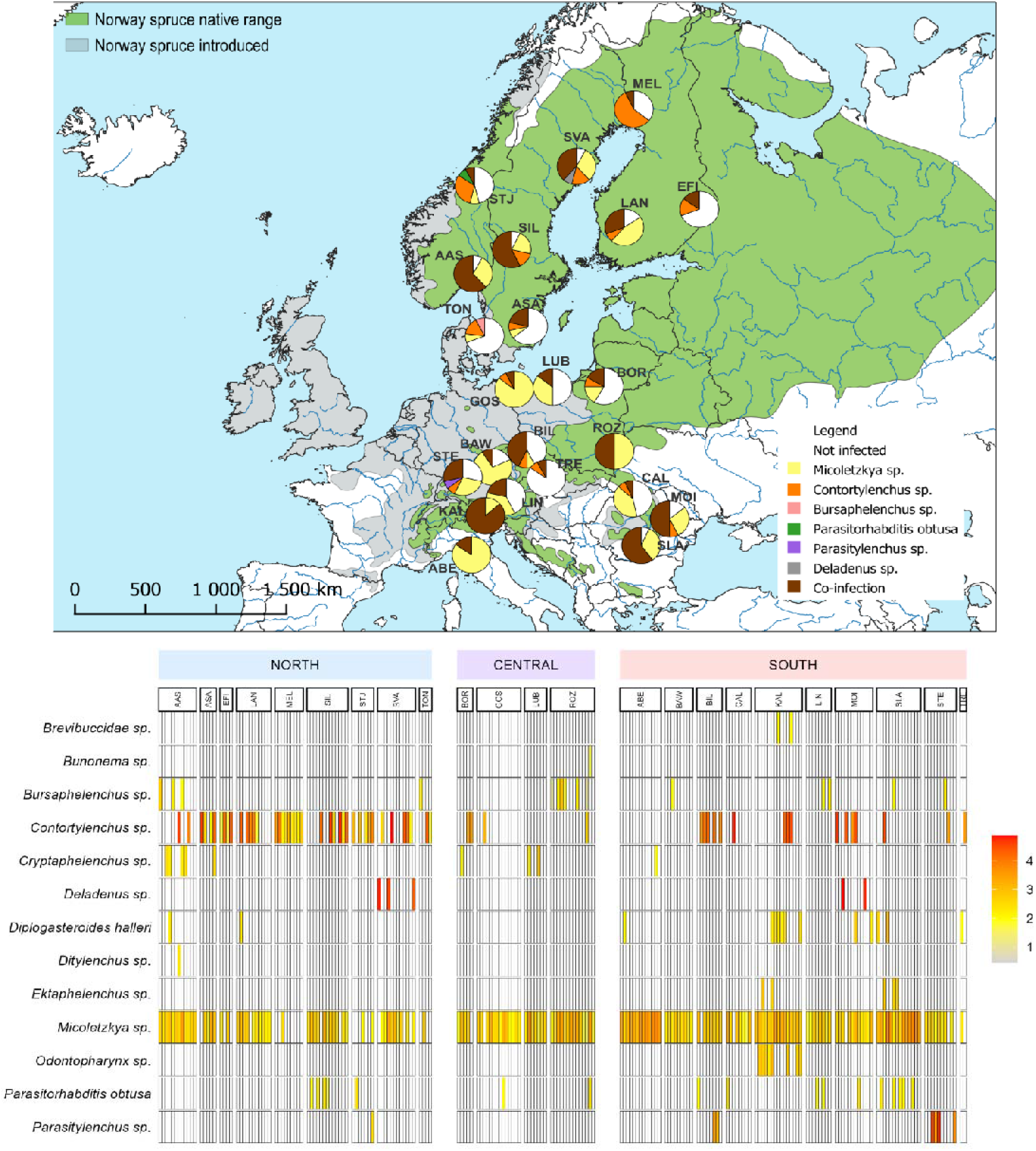
Geographic distribution of nematode species associated with *Ips typographus*. Top panel: map of sampling locations, with pie charts indicating the proportion of beetles with no nematode detection, detections of a single nematode species, or co-infections involving two or more nematode species that were seen in each population. Background shading indicated the native and introduced ranges of Norway spruce (*Picea abies*). Bottom panel: heatmap showing the prevalence of nematode species detected from metagenome-derived SSU rRNA sequences in beetle populations from Northern, Central and Southern regions. Rows correspond to nematode species and columns to individual beetles, grouped by population and geographic region. Cell colour represents log10(reads coverage + 1), allowing visualization of both high- and low-coverage detections; white cells indicate no detection. Nematode species are ordered alphabetically.

Other nematodes were detected less frequently. *Bursaphelenchus* sp. and *Parasitorhabditis obtusa* occurred in all three geographic groups, with no significant differences in prevalence among them (Fisher’s exact tests, p > 0.05). Several rare taxa were detected in only one group: *Ditylenchus* sp. in northern populations, *Bunonema* sp. in central populations, and *Ektaphelenchus* sp., *Odontopharynx* sp. and *Brevibuccidae* sp. in southern populations (Figure 2). With such low sample sizes, permutation tests found no significant geographic structuring for these taxa after correction for multiple testing (p > 0.05).

More than one nematode taxon was detected in 91 of the 312 beetles (29.2%). The maximum observed within an individual was four taxa, found in three beetles, including one from the central region and two from the southern region.

### Variable mitogenomes recovery success

With variable but often high nematode 18S rRNA sequence read coverage across beetle metagenomes, we attempted nematode mitochondrial genome reconstruction. GetOrganelle generated usable assemblies for 297 of 312 samples, yielding 362 total scaffold sequences. 305 scaffolds (363 - 18,243 bp; mean 14,528 bp; median 16,865 bp; IQR: 16,700 - 17,026 bp) were matched to the beetle mitochondrial genome with high confidence, whereas 51 scaffolds (820 - 14,603 bp; mean 8,983 bp; median 11,303 bp; IQR: 3,205 - 12,886 bp) confidently matched nematode mitogenomes. Due to variable success in protein-coding gene annotation, we restricted our downstream nematode phylogenetic analyses to the COX1 and COX2 genes. These genes were successfully recovered from 44 libraries, with mean lengths of 1505 bp and 679 bp, respectively.

Six scaffolds recovered from samples from Sweden (ASA124), Czech Republic (BIL34), and Germany (STE1, STE12, STE20, and STE24) lacked significant matches against the local reference database but were assigned to parasitoid wasps following searches against the full NCBI nucleotide database. These scaffolds ranged from 12,278 to 15,273 bp (mean: 13,696 bp; median: 14,006 bp) and yielded seven mitochondrial protein-coding genes: COX1, COX2, COX3, COB, NAD1, NAD4, and NAD5. Individual gene sequences ranged from 682 to 1,689 bp, with an overall mean length of 1,156 bp. COX1 comprised four distinct haplotypes and provided the strongest evidence for species identification, with all haplotypes matching a reference sequence of *Ropalophorus clavicornis* (Hymenoptera: Braconidae) at 99.1–99.4% nucleotide identity. This assignment is consistent with the recovery of these sequences from *Ips typographus*, whose adults are known hosts of this endoparasite (Wegensteiner et al., 2017).

### Limited genetic structure within nematode lineages

Phylogenetic reconstruction based on nematode SSU rRNA sequences recovered from *I. typographus* metagenomes revealed clear clustering by nematode taxon, with distinct lineages corresponding to the main genera detected in the dataset (Figure 3). As expected for SSU rRNA markers, within-taxon divergence was generally shallow and provided little information on the within-species variation.

**Figure 3.**
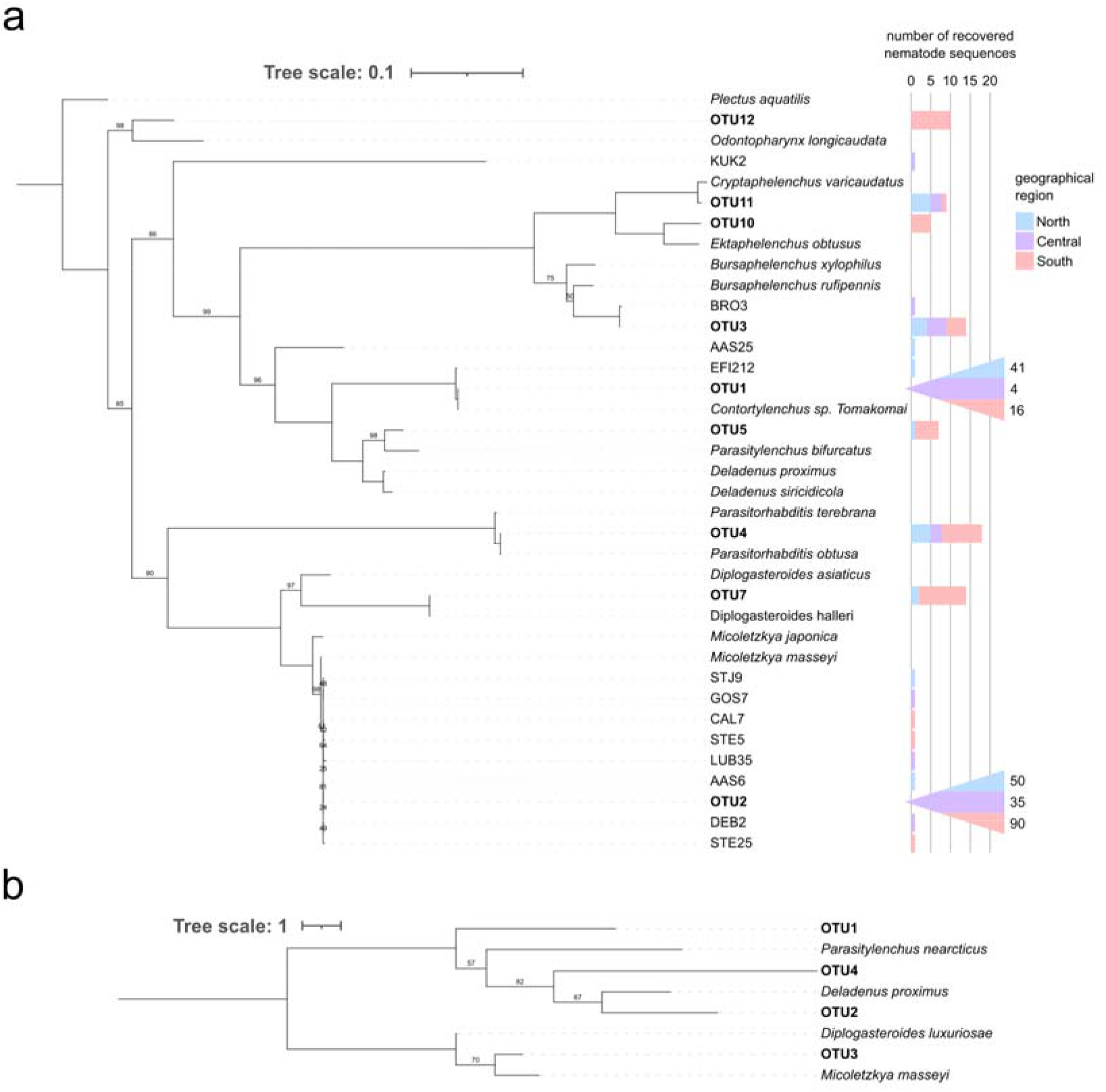
Maximum-likelihood phylogenies of off-target nematode sequences based on (a) SSU rRNA and (b) COX1. Recovered sequences were clustered separately for each marker into operational taxonomic units (OTUs) at 99% nucleotide identity. OTU numbers are marker specific. Italicized taxon names indicate public reference sequences, while other labels identify OTUs or sample identifiers for sequences that remained unmerged. The SSU tree was rooted using *Plectus aquatilis*, whereas the COX1 tree was midpoint rooted. Stacked bars in panel (a) show the number of recovered sequences associated with each tip, coloured by the geographical region of the corresponding host beetles. Branch labels indicate ultrafast bootstrap support (%) from 1,000 replicates; values of 100% are omitted. Scale bars indicate substitutions per site. Bars exceeding the displayed range are represented by triangles, with counts indicated by numbers.

Phylogenetic trees reconstructed based on COX1 and COX2 genes included just four of the nematode species detected in our data, and despite greater nucleotide sequence divergence, did not provide further insight due to the limited recovery success of the mitogenomes and the scarcity of available references in public databases.

## Discussion

Standard whole-genome resequencing analyses usually exclude reads that do not map to the reference genome, despite insights they can provide about other biologically relevant organisms present in the samples (Gouin et al., 2015). As more sequencing data become available, these reads offer opportunities to recover taxonomic marker sequences that can identify associated organisms and reveal how their occurrence varies across host populations. Even when the data do not always allow the reconstruction of high-quality genomes of these associated organisms, it can still help examine their distributions and population structure across host populations, as shown recently for the nematode *Contortylenchus reversus* in American Mountain pine beetle *Dendroctonus ponderosae* (Campbell et al., 2026; Campusano et al., 2026). While no high-quality genomic references are available so far for nematodes associated with *Ips* beetles, our study builds upon these approaches by examining several groups of organisms within a large resequencing dataset, showing how intermediate-coverage data generated to study the host can also be used to investigate its associations without additional sampling or sequencing.

The bacteria, fungi and mites detected in our datasets include taxa previously reported from *I. typographus*. For example, we have recovered four primary bacterial genera reported previously from bark beetles and their galleries: *Erwinia*, *Pseudomonas*, *Rahnella* and *Serratia* (Chakraborty et al., 2023; Peral-Aranega et al., 2023; Baños-Quintana et al., 2024). Among these, *Erwinia* and *Pseudomonas* have been reported to potentially contribute to beetle nutrition and protection against fungal pathogens (Peral-Aranega et al., 2023). Fungal detections mainly comprise yeasts, including *Wickerhamomyces bisporus*, which has previously been reported from the beetle gut (Veselská et al., 2023). We did not detect known ophiostomatoid associates such as *Endoconidiophora polonica* and *Ophiostoma bicolor*, although a recent preprint reports that adult *I. typographus* carry fungal symbionts in mandibular mycetangia (Lehenberger et al., 2026). Our DNA extraction protocol was optimized for beetle tissue, and the subsequent screen relied on the recovery and taxonomic assignment of SSU rRNA sequences, both of which may have limited our ability to detect these fungi. Similarly, mites of the genus *Trichouropoda* are known to be carried by adult *I. typographus* (Toma et al., 2025). Comparatively less has been known about the communities of nematodes associated with *Ips* beetles. Our detection of nematodes in more than two-thirds of individuals, and the high proportion of DNA that they often comprise, highlight the need to examine this group alongside the other organisms associated with *I. typographus*.

Our analysis revealed thirteen nematode taxa associated with spruce bark beetle populations across Europe. Several genera recovered here, including *Micoletzkya*, *Bursaphelenchus*, *Cryptaphelenchus*, *Ektaphelenchus*, *Contortylenchus* and *Parasitylenchus*, have previously been recorded from *I. typographus* (Grucmanová & Holuša, 2013) and related species. For example, surveys of the closely related bark beetle species *I. cembrae* have recovered *Micoletzkya* from the beetle surface, while genera *Contortylenchus*, *Parasitylenchus* and *Cryptaphelenchus* are known to inhabit beetle body cavities (Grucmanová et al., 2016). Similarly, *Bursaphelenchus*, *Micoletzkya* and *Contortylenchus* have been recorded in the nematode fauna of *Dryocoetes uniseriatus* (Shimizu et al., 2013), a beetle species belonging to the same subfamily (Scolytinae) as *I. typographus*. The nematode taxa detected here differ in the nature of their relationships with their beetle hosts. *Micoletzkya*, for example, uses beetles for transport and feeds on other organisms in their galleries, including other nematodes, whereas *Contortylenchus* and *Parasitylenchus* develop within beetle body cavities as parasites (Susoy & Herrmann, 2014; Grucmanová et al., 2016).

Other nematode detections are harder to relate to established associations with *I. typographus*. *Deladenus*, *Bunonema*, *Odontopharynx* and *Brevibuccidae* are not included among the taxa listed for central European *Ips* species by Grucmanová and Holuša (2013). *Bunonema* has been reported among associates of other bark beetle genera, while *Deladenus* includes species with fungal-feeding and insect-parasitic stages (Carta et al., 2010; Kosaka & Ogura, 1993). For these less documented detections, systematic examination of beetles and their galleries would help distinguish organisms carried by the beetle from those encountered in the same environment.

These nematode genera differ in their distribution across genetically differentiated host beetle clades. However, besides host genetic structure, nematode occurrence is also likely to be influenced by the environmental conditions experienced by different life stages. The contrasting geographic distributions of *Micoletzkya* and *Contortylenchus* could reflect such differences. For nematodes transported between galleries, occurrence may depend on both accesses to beetle carriers and conditions for feeding and reproduction in the gallery. For internal parasites, it also depends on successful infection and development within the beetle. The survival and performance of life stages is likely to depend on geographically structured drivers such as temperatures. Previous work shows that these associations can vary over space and time: nematode infestation differed among locations and generations of *I. cembrae*, while seasonal changes in nematode associations were recorded in *D. uniseriatus* (Grucmanová et al., 2016; Shimizu et al., 2013). The geographic patterns that we report are likely explained by various combinations of local environmental conditions, sampling time and host-related factors.

The higher prevalence of *Contortylenchus* in northern populations is particularly interesting given the parasitic nematodes’ reported effects on beetle reproduction. In *I. typographus*, nematodes in the hemocoel have been associated with fewer eggs and shorter maternal galleries, whereas intestinal nematodes did not show the same relationship with reproduction (Resnerová et al., 2022). In another wood-boring beetle, *Monochamus alternatus*, *C. genitalicola* parasitises the reproductive organs and has been considered a potential biological control agent because of its effects on reproduction (Kosaka & Ogura, 1993; Shimizu et al., 2013). The significance of geographic differences in *Contortylenchus* prevalence to beetle reproduction and outbreak potential remain to be investigated. Geographic variation has also been examined at the genetic level: Campusano et al. (2026) recovered *C. reversus* from *Dendroctonus ponderosae* sequencing data and found that its population structure broadly corresponded to that of the beetle. Together with our observations of geographic differences in prevalence, this raises the question of how host population history and local conditions contribute to the distribution of these parasites.

In multiple cases, different nematode taxa co-occurred within the same beetles. Co-infections by different nematode specie have been documented in other bark beetles, sometimes without clear evidence of competition (Shimizu et al., 2013). Co-occurrence can therefore reflect shared exposure or the use of different parts of the beetle, rather than a direct interaction between nematodes. Nevertheless, the predatory habits of *Micoletzkya* could affect the survival or abundance of other nematodes in galleries, or of their juvenile stages as they attempt to colonize beetles’ body cavities (Susoy & Herrmann, 2014). Whether such potential interactions might influence beetle outbreaks remains unresolved; for some nematode taxa, no significant trends during outbreaks were seen (Holuša & Lukášová, 2017). Repeated sampling would be needed to establish whether the combinations detected here persist across beetle generations or change with population size.

In our survey, metagenome-reconstructed SSU rRNA sequences distinguished the main nematode lineages but provided limited insight into within-lineage genetic variation, a critical aspect of host-parasite community comparisons. Mitochondrial and nuclear-encoded protein-coding genes can offer substantially greater phylogenetic resolution but may not be easy to retrieve from resequencing short-read datasets, due to mitogenomes’ low GC contents (Campbell et al., 2026), parasites’ nuclear genes’ low read coverage, and the lack of suitable references for sequence identification and comparison. In our study, the retrieval success of nematodes’ mitogenomes, or even mitochondrial genes such as cox1 and cox2, was surprisingly low, and references for comparisons scarce in public databases. Expanding the set of publicly available reference sequences for bark beetle-associated nematodes would help improve taxonomic identification and phylogenetic placement of sequences recovered from host resequencing data.

Despite these limitations, our study fills gaps in our knowledge of nematodes associated with *I. typographus*, particularly their geographic distribution and co-occurrence in individual beetles. Our results show that whole-genome resequencing datasets can provide valuable information on several nematode taxa and diverse other organisms simultaneously across a broad geographic range, offering a basis for further studies of their relationships with spruce bark beetles.

## Conclusions and future directions

Genomic datasets generated to study *I. typographus* genomes contain abundant information about its biological associations that would otherwise be discarded during standard analyses. Using such data, we document surprisingly high nematodes’ prevalence across beetle populations, alongside variation in nematode communities among beetle populations and the co-occurrence of different taxa within individuals. On the other hand, the phylogenetic resolution offered by readily retrievable rRNA sequences is limited, and other regions of the genome are substantially harder to reconstruct. Nevertheless, our data provide a strong foundation for future studies of the factors shaping these patterns and their consequences for beetle survival and reproduction. Future studies that combine targeted molecular screening across populations, timepoints, and life stages, alongside measurements of beetle performance, will help determine whether associated organisms contribute to variation in spruce bark beetle population dynamics and outbreak development.

## Acknowledgements

This project was supported by the Polish National Science Centre grants 2018/30/E/NZ8/00105 to KNB and 2021/43/B/NZ8/03376 to PŁ. We also acknowledge Bernardo Flores Antunes for his help with graphic visualisation of the results.

## Notes

### Competing Interest Statement

The authors have declared no competing interest.

