## Supplementary Information for "High prevalence and diversity of nematodes associated with Europe’s most destructive forest pest, revealed by off-target genomic data"

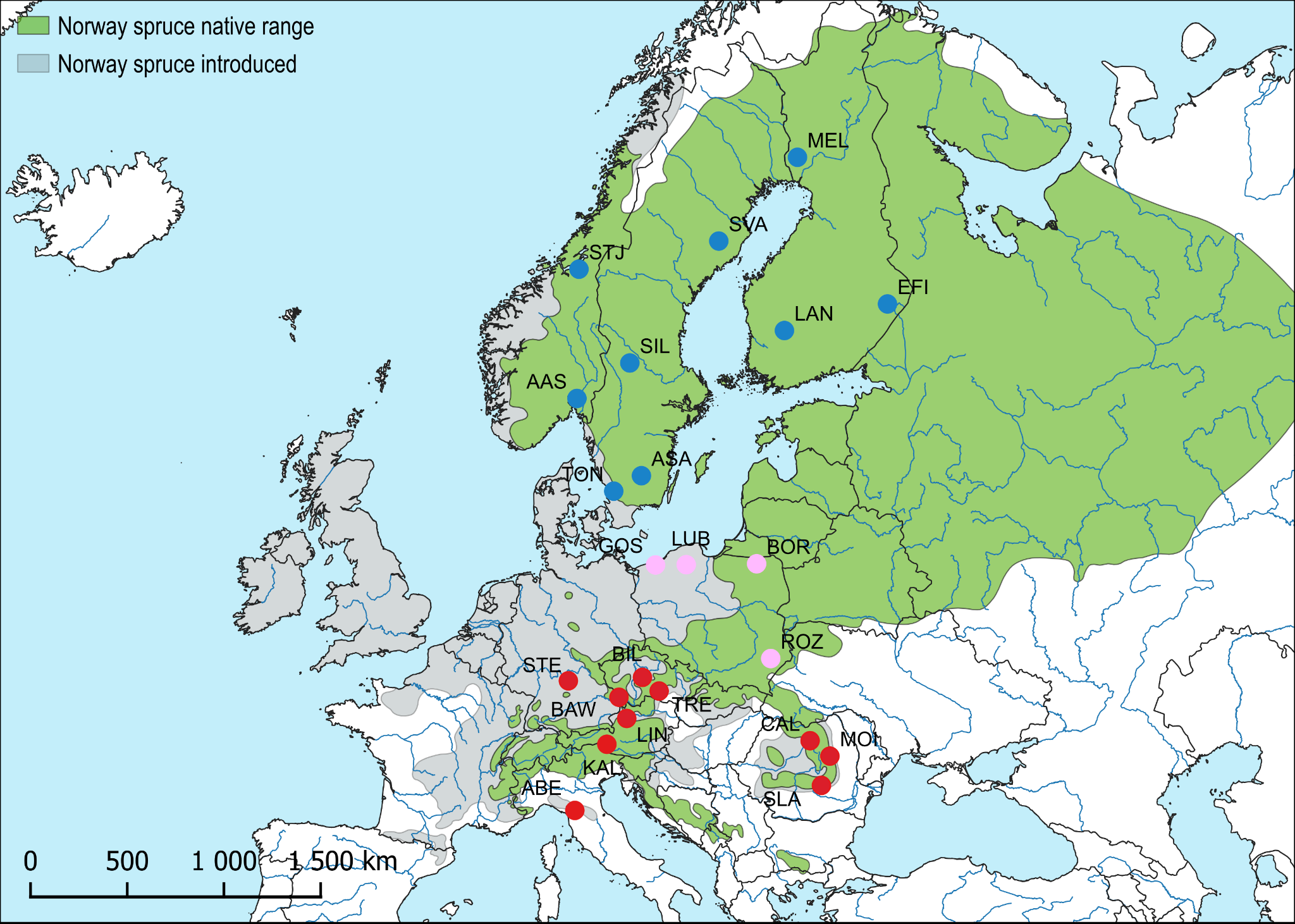

**Figure S1.** Sampling sites and assignment to *Ips typographus* genetic groups. Blue - populations assigned to the northern group, red - southern group, pink - central group (populations being highly admixed between southern and northern *I. typographus* genetic groups). Population labels correspond to Population ID in Table S1.

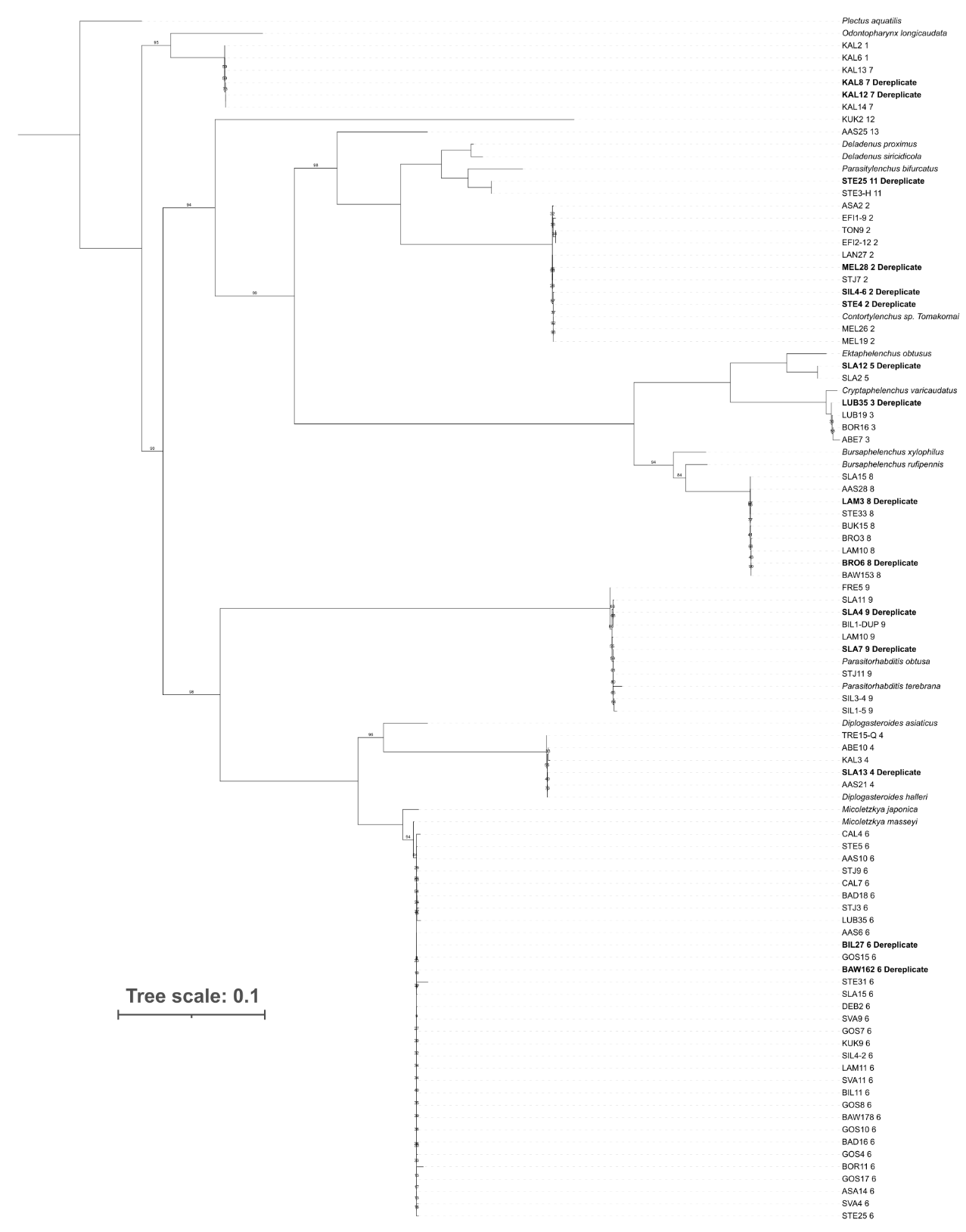

**Figure S2.** Nematode SSU rRNA phylogeny generated using dereplicated sequences obtained from PhyloFlash. When multiple sequences were identical, only one sample name was kept. Bootstrap labels for branches with 100% support were omitted.

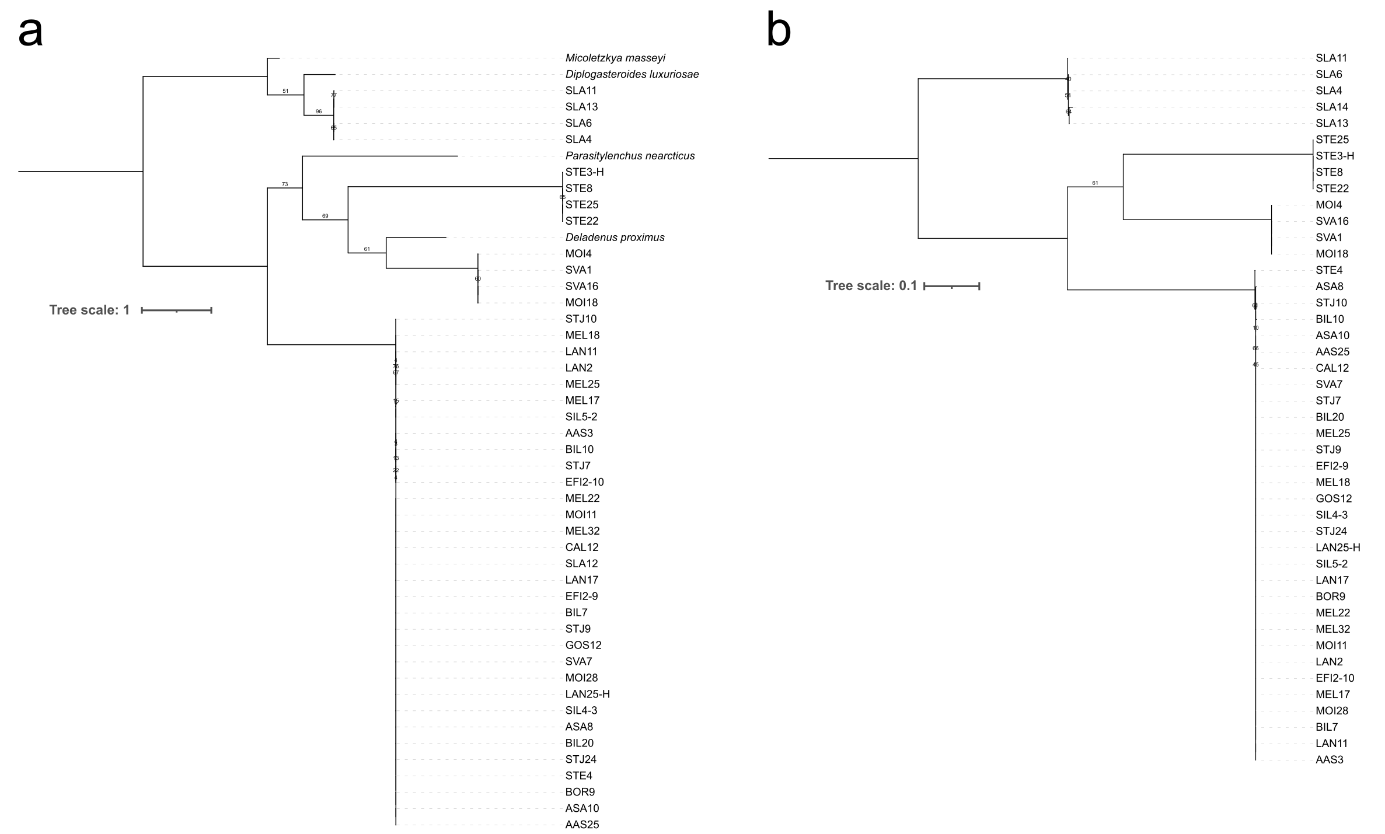

**Figure S3.** Nematode phylogenies generated using a) COX1 and b) COX2 genes obtained from GetOrganelle. Both phylogenies were rooted at the midpoint. Bootstrap labels were omitted for branches with 100% support.

**Table S1.** *Ips typographus* sampling information. ‘Group’ indicates in what part of the species’ distribution area beetles were collected: northern (N), southern (S) and central (C). ‘N’ indicates the number of individuals collected per population.

| Population ID | Locality ID | N | Population name | Latitude | Longitude | Country | Group |
| --- | --- | --- | --- | --- | --- | --- | --- |
| ABE | ABE | 13 | Abetone | 44.148 | 10.663 | Italy | S |
| KAL | KAL | 15 | Kals | 47.012 | 12.646 | Austria | S |
| LIN | LAM, FRE, BAD | 14 | Linz | 48.092 | 13.874 | Austria | S |
| BAW | BAW | 11 | Bavarian Forest | 48.960 | 13.395 | Germany | S |
| TRE | TRE | 13 | Trebic | 49.212 | 15.879 | Czech | S |
| STE | STE | 14 | Steigerwald | 49.622 | 10.263 | Germany | S |
| BIL | BIL | 14 | Bilkovice | 49.761 | 14.848 | Czech | S |
| SLA | SLA | 15 | Slanic | 45.250 | 25.925 | Romania | S |
| MOI | MOI | 14 | Moinesti | 46.510 | 26.449 | Romania | S |
| CAL | CAL | 15 | Calimani | 47.163 | 25.221 | Romania | S |
| ROZ | BRO, DEB, KUK, BUK | 14 | Roztocze | 50.508 | 22.786 | Poland | C |
| GOS | GOS | 14 | Goscinno | 54.047 | 15.657 | Poland | C |
| LUB | LUB | 14 | Lubaszki | 54.057 | 17.556 | Poland | C |
| BOR | BOR | 12 | Borki | 54.090 | 21.912 | Poland | C |
| TON | TON | 13 | Tönnersjö | 56.643 | 13.070 | Sweden | N |
| ASA | ASA | 14 | Asa | 57.165 | 14.783 | Sweden | N |
| AAS | AAS | 13 | As | 59.667 | 10.793 | Norway | N |
| SIL | SIL | 14 | Siljanfors | 60.757 | 14.066 | Sweden | N |
| LAN | LAN | 13 | Länsi | 61.723 | 23.633 | Finland | N |
| EFI | EFI | 13 | Eastern Finland | 62.492 | 30.010 | Finland | N |
| STJ | STJ | 13 | Stjordal | 63.469 | 10.918 | Norway | N |
| SVA | SVA | 13 | Svartberget | 64.236 | 19.570 | Sweden | N |
| MEL | MEL | 14 | Mellakoski | 66.399 | 24.440 | Finland | N |

**Table S2.** Selected mitochondrial genomes used as GetOrganelle references. Nematode references were the closest available mitochondrial genomes at the indicated taxonomic level (genus, family or order) for the taxa detected; the *Ips typographus* mitogenome was included as the host reference. Accession numbers link to the corresponding NCBI Nucleotide records.

| **Taxon** | **Accession** | **Mitochondrial record** | **Reference level** | **Reference** |
| --- | --- | --- | --- | --- |
| *Allodiplogaster sudhausi** | [KT355738.1](https://www.ncbi.nlm.nih.gov/nuccore/KT355738.1) | Complete genome, 16,005 bp | Order (Rhabditida) | Kim T, Kim J, Nadler SA, Park JK (2016) *Curr. Genet.* 62: 391–403. doi:10.1007/s00294-015-0536-4 |
| *Bunonema* sp.** | [MT920392.1](https://www.ncbi.nlm.nih.gov/nuccore/MT920392.1) | Mitochondrial sequence, 16,601 bp | Genus | Casasa S, Biddle JF, Koutsovoulos GD, Ragsdale EJ (2021) *Mol. Biol. Evol.* 38: 331–343. doi:10.1093/molbev/msaa235 |
| *Bursaphelenchus sinensis* | [PV492208.1](https://www.ncbi.nlm.nih.gov/nuccore/PV492208.1) | Complete genome, 18,386 bp | Family (Aphelenchoididae) | Huang XJ, Li HM, Qing X (2025) Unpublished; GenBank direct submission |
| *Bursaphelenchus xylophilus* isolate BxPt19SCD | [JQ514068.1](https://www.ncbi.nlm.nih.gov/nuccore/JQ514068.1) | Complete genome, 12,945 bp | Genus | Moreira C, Pereira F, Fonseca L, van Asch B, Mota M, Abrantes I, Amorim A (2012) Unpublished; GenBank direct submission |
| *Diplogasteroides magnus*** | [MT920393.1](https://www.ncbi.nlm.nih.gov/nuccore/MT920393.1) | Mitochondrial sequence, 14,844 bp | Genus | Casasa S, Biddle JF, Koutsovoulos GD, Ragsdale EJ (2021) *Mol. Biol. Evol.* 38: 331–343. doi:10.1093/molbev/msaa235 |
| *Pratylenchus vulnus* | [GQ332425.1](https://www.ncbi.nlm.nih.gov/nuccore/GQ332425.1) | Complete genome, 21,656 bp | Order (Tylenchida) | Sultana T, Kim J, Lee SH, Han H, Kim S, Min GS, Nadler SA, Park JK (2013) *BMC Evol. Biol.* 13: 12. doi:10.1186/1471-2148-13-12 |
| *Ips typographus* isolate JGIT1 (host beetle) | [OK110249.1](https://www.ncbi.nlm.nih.gov/nuccore/OK110249.1) | Partial genome, 16,705 bp | Host | Du H (2021) Unpublished; GenBank direct submission |

* Published as *Koerneria sudhausi*; the record is now listed under *Allodiplogaster sudhausi*. ** GenBank “UNVERIFIED” record: sequence and annotation have not been verified by GenBank staff.

**Table S3.** Selected reference mitochondrial COI sequences of nematodes retrieved from GenBank/ENA (accessed 29 September 2026) for available genera detected in this study. Accession numbers link to the corresponding NCBI Nucleotide records. Locality and publication title are given as stated in the sequence record; “Not stated” indicates that the record lacks this information.

| **Organism** | **Accession** | **Locality** | **Publication title** |
| --- | --- | --- | --- |
| *Micoletzkya japonica* | [JX163957.1](https://www.ncbi.nlm.nih.gov/nuccore/JX163957.1) | Japan | Description of bark-beetle associated nematodes Micoletzkya masseyi n. sp. and M. japonica n. sp. (Nematoda: Diplogastridae) |
| *Micoletzkya masseyi* | [JX163958.1](https://www.ncbi.nlm.nih.gov/nuccore/JX163958.1) | Canada | Description of bark-beetle associated nematodes Micoletzkya masseyi n. sp. and M. japonica n. sp. (Nematoda: Diplogastridae) |
| *Diplogasteroides asiaticus* | [LC027676.1](https://www.ncbi.nlm.nih.gov/nuccore/LC027676.1) | Japan: Ibaraki, Kasumigaura, Chiyoda Experimental Station of the Forestry and Forest Products Research Institute | Diplogasteroides asiaticus n. sp. is Associated with Monochamus alternatus in Japan; Allodiplogaster josephi n. sp. and A. seani n. sp. (Nematoda: Diplogastridae), associates of soil-dwelling bees in the eastern United States |
| *Diplogasteroides luxuriosae* | [LC099976.1](https://www.ncbi.nlm.nih.gov/nuccore/LC099976.1) | Japan: Ibaraki, Tsukuba, Forestry and Forest Products Research Institute | Diplogasteroides luxuriosae n. sp. associated with Acalolepta luxuriosa (Coleoptera: Cerambycidae) from Japan |
| *Parasitorhabditis obtusa* | [PQ585736.1](https://www.ncbi.nlm.nih.gov/nuccore/PQ585736.1) | South Korea | Description of a New Cryptic Rhabditid, Parasitorhabditis paraterebrana n. sp. (Nematoda: Rhabditidae), with Remarks on Two Known Species from Korea |
| *Parasitorhabditis terebrana* | [OQ281743.1](https://www.ncbi.nlm.nih.gov/nuccore/OQ281743.1) | South Korea | Morphological and Molecular Characterization of Diplogasteroides sp., a Cryptic Population of the Haslacheri Group (Diplogastridae), and Parasitorhabditis terebranus (Rhabditidae) from Korea |
| *Bursaphelenchus rufipennis* | [AB368528.1](https://www.ncbi.nlm.nih.gov/nuccore/AB368528.1) | Canada: Fredericton | Bursaphelenchus rufipennis n. sp. (Nematoda: Parasitaphelenchinae) and redescription of Ektaphelenchus obtusus (Nematoda: Ektaphelenchinae), associates from nematangia on the hind wings of Dendroctonus rufipennis (Coleoptera: Scolytidae) |
| *Bursaphelenchus xylophilus* | [AP017463.1](https://www.ncbi.nlm.nih.gov/nuccore/AP017463.1) | Japan | Complete mitochondrial genomes of 50 helminths species |
| *Cryptaphelenchus* sp. o XM-2023 | [OR223593.1](https://www.ncbi.nlm.nih.gov/nuccore/OR223593.1) | China | Unveiling novel and known Cryptaphelenchus species from China and USA (Unpublished) |
| *Parasitylenchus nearcticus* | [AF519208.1](https://www.ncbi.nlm.nih.gov/nuccore/AF519208.1) | Not stated | Infection success in novel hosts: an experimental and phylogenetic study of Drosophila-parasitic nematodes |
| *Ektaphelenchus obtusus* | [AB368531.1](https://www.ncbi.nlm.nih.gov/nuccore/AB368531.1) | Canada: Fredericton | Bursaphelenchus rufipennis n. sp. (Nematoda: Parasitaphelenchinae) and redescription of Ektaphelenchus obtusus (Nematoda: Ektaphelenchinae), associates from nematangia on the hind wings of Dendroctonus rufipennis (Coleoptera: Scolytidae) |
| *Ektaphelenchus joyceae* | [MT712703.1](https://www.ncbi.nlm.nih.gov/nuccore/MT712703.1) | China: Liaoning | First Report of Ektaphelenchus joyceae (Rhabditida: Ektaphelenchinae) on Korean Pine Pinus koraiensis and Black Pine Pinus thunbergii in China |
| *Deladenus siricidicola* | [EU545474.1](https://www.ncbi.nlm.nih.gov/nuccore/EU545474.1) | Canada: Ontario | Characterization of Deladenus siricidicola associated with Sirex noctilio in Ontario, Canada |
| *Deladenus proximus* | [KJ193977.1](https://www.ncbi.nlm.nih.gov/nuccore/KJ193977.1) | Not stated | Distribution and genetic structure of Deladenus proximus, a nematode parasite of the woodwasp Sirex nigricornis in the eastern United States |

**Table S4.** Concordance of nematode detections between original and duplicate extractions from the same individuals, based on the two most abundant taxa (*Contortylenchus* sp. and *Micoletzkya sp.*). Each pair of values corresponds to two independent extractions and sequencing runs from the same individual, shown in the format original/duplicate (i.e. SVA8 (original)/SVA8-DUP (duplicate)). Concordance was defined as both extractions giving the same result (either presence or absence), which occurred in 83% of cases.

|  | Samples | | | | | | | | |
| --- | --- | --- | --- | --- | --- | --- | --- | --- | --- |
| Nematode taxa | SVA8/  SVA8-DUP | STJ1/  STJ1-DUP | STE4/  STE4-DUP | SIL1-4/  SIL1-4-DUP | LUB1  /LUB1-DUP | LAN1  /LAN1-DUP | EFI1-7/  EFI1-7-DUP | BRO3/  BRO3-DUP | AAS25/  AAS25-DUP |
| *Contortylenchus sp.* | 0/1 | 1/0 | 1/1 | 0/0 | 0/0 | 0/0 | 0/0 | 0/0 | 1/1 |
| *Micoletzkya sp.* | 0/0 | 0/0 | 0/1 | 1/1 | 0/0 | 1/1 | 0/0 | 1/1 | 1/1 |

**Table S5.** Taxonomy and published ecological associations of the 13 nematode taxa detected by assembled SSU rRNA sequences. Geographic groups refer to detections in this study (N, northern Europe; C, central; S, southern Europe), not the entire geographic ranges of the nematodes. Published associations refer to the named genus or a specified species and should not be interpreted as direct observations of life stage or location in our beetles.

| **Detected taxon** | **Order; family** | **Groups** | **Location or ecological relationship** | **Taxonomic notes** | **References*** |
| --- | --- | --- | --- | --- | --- |
| *Micoletzkya* sp. | Diplogastrida; Diplogastridae | N, C, S | Phoretic on adult bark beetles (often under elytra); develops in galleries and feeds on associated organisms, including nematodes. Not established as an *Ips* parasite. | — | [1], [2] |
| *Diplogasteroides halleri* | Diplogastrida; Diplogasteroididae | N, S | Reported on external body surfaces of *Ips*; phoretic/gallery associate. | Also listed historically as *Fuchsnema halleri*; taxonomic placement needs checking. | [1] |
| *Parasitorhabditis obtusa* | Rhabditida; Rhabditidae | N, C, S | Intestinal associate/endoparasite reported from *I. typographus*; lesions in gut epithelium reported in experimental work, though effects vary by host and study. | — | [1], [3] |
| *Odontopharynx* sp. | Diplogasterida; Odontopharyngidae | S | Predatory, free-living nematode at genus level; direct parasitism or a specific transport site in *I. typographus* is undocumented. | — | [4] |
| *Contortylenchus* sp. | Tylenchida; Allantonematidae | N, C, S | Insect endoparasite; *C. diplogaster* is recorded in the hemocoel of *I. typographus* in central Europe and Bulgaria. | — | [1] |
| *Bursaphelenchus* sp. | Tylenchida; Aphelenchoididae | N, C, S | Typically, phoretic on bark beetles and reproduces in wood/galleries; some species feed on fungi. *B. eidmanni* is recorded on *I. typographus*. | Family also placed in Parasitaphelenchidae in older sources. | [1] |
| *Cryptaphelenchus* sp. | Tylenchida; Aphelenchoididae | N, C, S | Reported both as phoretic on beetles and within intestine/Malpighian tubules in *Ips*. The association in our beetles is unresolved. | — | [1] |
| *Parasitylenchus* sp. | Tylenchida; Parasitylenchidae | N, S | Insect endoparasite; *P. dispar* has been recorded in the hemocoel of *I. typographus*. | — | [1] |
| *Ektaphelenchus* sp. | Tylenchida; Ektaphelenchidae | S | *Ips*-associated genus with phoretic stages under elytra/on wings, sometimes in nematangia; *E. typographi* is also reported from beetle larvae. | — | [1] |
| *Deladenus* sp. | Tylenchida; Neotylenchidae | N, S | Some congeners alternate between fungal-feeding and insect-parasitic phases, especially in woodwasps. No evidence here establishes parasitism of *I. typographus*. | PhyloFlash originally labelled these sequences *Urocerus augur*; BLAST supported *Deladenus* sp. | [5] |
| *Ditylenchus* sp. | Tylenchida; Anguinidae | N | Ecology unresolved at genus level. | Historical *Ditylenchus major* from Ips is now treated as *Neoditylenchus major*; our sequence is not assigned to that species. | [1] |
| *Bunonema* sp. | Rhabditida; Bunonematidae | C | Free-living bacterivore; phoresy has been reported for congeners on other insects. Direct relationship with *I. typographus* is unresolved. | — | [6] |
| *Brevibuccidae* sp. | Rhabditida; Brevibuccidae | S | Family includes free-living forms found in bark-beetle tunnels; family-level detection cannot establish feeding mode or parasitism in this beetle. | — | [7] |

***References**

**Other Materials**. Detailed table of phyloFlash SSU rRNA reads detection.

https://drive.google.com/drive/folders/1jYdFSl8C2NRjFrELnvdR_jEGC4DIGc42?usp=drive_link
